# Compensation of Hyperexcitability with Simulation-Based Inference

**DOI:** 10.1101/2025.01.07.631838

**Authors:** Daniel Müller-Komorowska, Tomoki Fukai

## Abstract

The activity of healthy neuronal networks is tightly regulated, and a shift towards hyperexcitability can cause various problems, such as epilepsies, memory deficits, and motor disorders. Numerous cellular, synaptic, and intrinsic mechanisms of hyperexcitability and compensatory mechanisms to restore healthy activity have been proposed. However, quantifying multiple compensatory mechanisms and their dependence on specific pathophysiological mechanisms has proven challenging, even in computational models. We use simulation-based inference to quantify the interactions of putative compensatory mechanisms in a spiking neuronal network model. Various parameters of the model can compensate for changes in other parameters to maintain baseline activity, and we estimated their compensatory potential. Furthermore, specific causes of hyperexcitability - interneuron loss, excitatory recurrent synapses, and principal cell depolarization - have distinct compensatory mechanisms that can restore normal excitability. Our results show that spiking neuronal network simulators could generate hypotheses about the mechanisms of pathophysiological network mechanisms.

## 1 Introduction

Neuronal networks operate within narrow ranges of average activity. A shift outside that range is a key aspect of many brain disorders. Epilepsy is an extreme case where hyperactivity of a neuronal network spreads across the brain, causing seizures (Devinsky et al., 2018). Alzheimer’s disease shows local hyperexcitability at the early stage, increased bursting, and elevated intracellular calcium levels (Anastacio et al., 2022; Mittag et al., 2023). Although this kind of hyperexcitability rarely causes seizures, there are similarities between local patterns of hyperactivity in epilepsy and Alzheimer’s disease (Kamondi et al., 2024). In schizophrenia, the imbalance of excitation and inhibition shifts cortical networks toward hyperactivity (Liu et al., 2021).

Hyperexcitability can result from changes at the cellular, synaptic, or molecular levels. At the cellular level, animal models of epilepsy show loss of interneurons (Huusko et al., 2015), and interneuron ablation leads to hyperexcitability (Martin & Sloviter, 2001). The role of interneurons in network excitability also led to the idea that grafting of new interneurons could prevent seizures (Hunt & Baraban, 2015).

At the synaptic level, increased excitatory connectivity can lead to hyperexcitability. Granule cells in the healthy dentate gyrus lack recurrent excitatory connections, whereas epileptic tissue exhibits aberrant recurrent excitation due to mossy fiber sprouting (Scharfman et al., 2003). Besides the recurrent connections, synaptic inputs from the medial perforant path to the dentate gyrus are also strengthened (Janz et al., 2017).

At the molecular level, mutations that affect ion channel functions can cause genetic epilepsy, and acquired forms of epilepsy are often accompanied by changes in ion channel expression or localization (Beck & Yaari, 2008; Lerche et al., 2013). But changes in ion channel expression can also restore healthy activity after injury (Yim et al., 2015). This shows that some ion channels can compensate for perturbations of other ion channels. Different ion channels performing the same function are an example of degeneracy (Marom & Marder, 2023). Degeneracy could be an important property of neuronal systems because it suggests that the same disorder can have different causes and that multiple compensatory mechanisms could restore healthy function (Stöber et al., 2023).

Studying the interactions of multiple, degenerate compensatory mechanisms across scales is extremely difficult in biological systems. But computer simulations provide a way to explore plausible mechanisms efficiently. A simulator produces output given parameters, which can be used directly to observe the effects of changing those parameters, similar to an experimental manipulation. But to identify degenerate compensatory mechanisms, it is more important to solve the inverse problem; to find the parameters that are likely to produce a given output.

Simulation-based inference (SBI) is widely used across scientific disciplines to solve the inverse problem of simulators (Cranmer et al., 2020). While there are different methods for performing SBI, they all use parameter-output samples for inference. For example, approximate Bayesian computation (ABC) infers the parameter distribution by discarding parameter samples that yield output very different from the target output. ABC is widely used, but discarding samples is inefficient, which makes it infeasible for biologically plausible simulators.

Neural posterior estimation (NPE) was recently developed as a more sampling efficient SBI method (Papamakarios & Murray, 2018). It is more sampling efficient because it uses a simulator’s output to train a neuronal density estimator to estimate the parameter distribution. Like other SBI methods, NPE works on any kind of simulator and is widely used across scientific fields. In neuroscience, it was used on neuronal models such as the stomatogastric ganglion system (Gonçalves et al., 2020) and on field recordings from various mouse brain areas (Gao et al., 2024).

We use NPE to identify putative compensatory mechanisms that avoid hyperexcitability in a spiking microcircuit simulator with two interneuron populations. We specifically investigate three well-known pathophysiological issues: interneuron (IN) loss, excitatory recurrent sprouting, and principal cell (PC) depolarization. We find that various parameters related to these three conditions have compensatory potential. Which parameters are most effective depends on the specific pathophysiology. IN loss is best compensated by increasing IN → PC connectivity. Sprouting is compensated by a set of PC intrinsic properties and PC → PC synaptic weight. Intrinsic depolarization is compensated best by increasing the PC spiking threshold. Our results show that identifying the specific pathophysiology could be used to estimate compensatory mechanisms.

## 2 Results

To study the mechanisms of hyperexcitability, we implemented a phenomenological spiking neuronal microcircuit simulator (Figure 1 A, Methods Section 4.1). The model has 32 free parameters describing intrinsic neuronal properties, synaptic strengths, connection probabilities, and interneuron numbers. The model’s outputs are 7 summary statistics that describe the average activity, firing regularity and oscillation strengths. To explore the model’s high dimensional parameter space, we defined a broad, uniform prior distribution of the input parameters (Table 1 & Table 2) and simulated 2,620,000 samples from it. The simulator output varied widely, from sparse, irregular spiking to hypersynchronous bursts (Figure 1 B & C).

**Figure 1:**
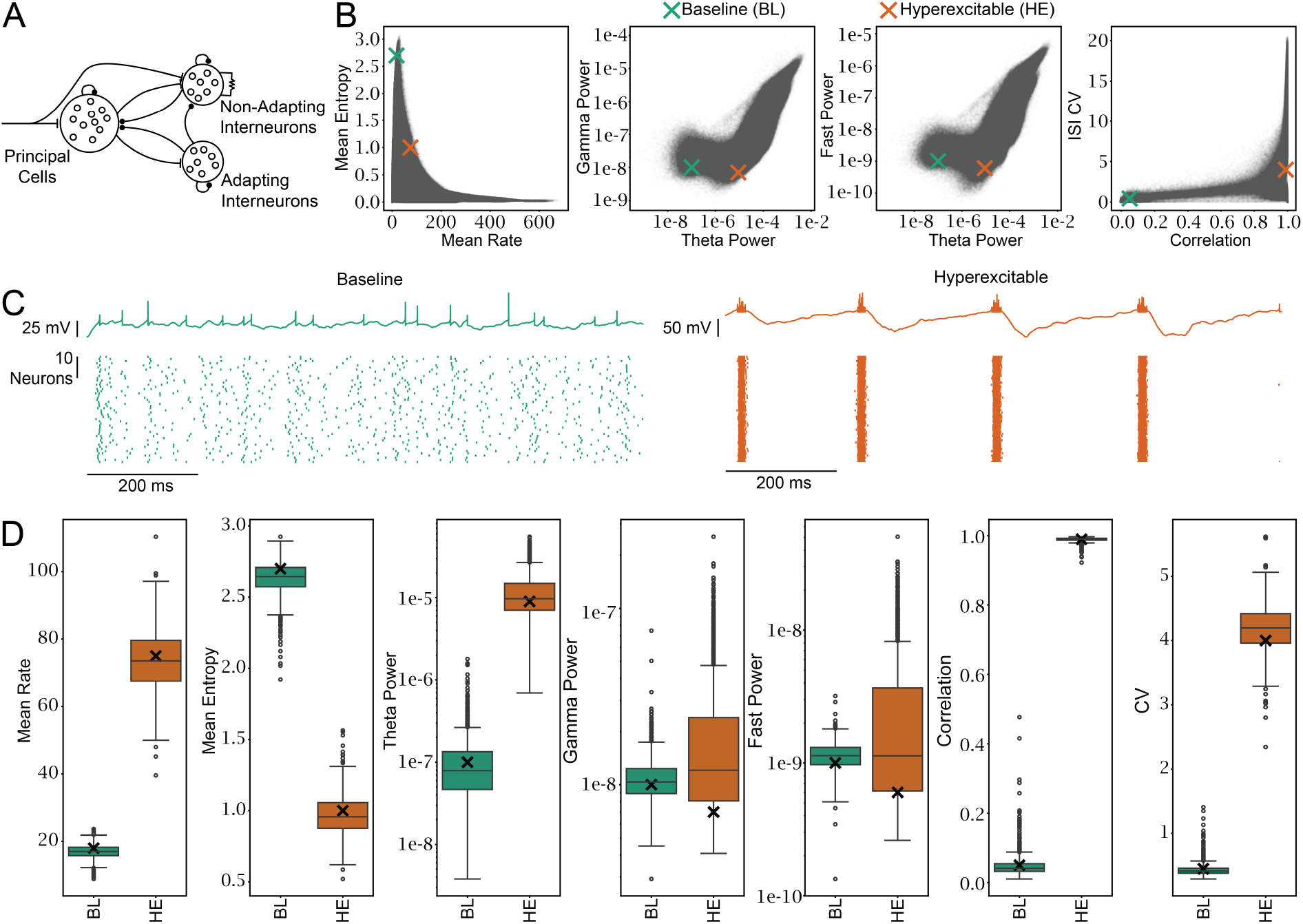
Sequential NPE finds distributions of parameters that produce different levels of excitability. **A)** An illustration of the spiking neuronal network simulator. See methods Section 4.1 for details. **B)** Simulator outcome from 1122586 simulations. Simulations with zero PC spikes or undefined CV are not shown. The parameters were drawn from the prior distribution (see Table 1 & Table 2). The cross shows the two target outcomes the NPE was conditioned on. **C)** Shows the result of simulating the MAP parameters from *p*(*θ* |*x*_BL_) and *p*(*θ* |*x*_HE_). Top: Voltage trace of a single PC. Bottom: spike raster plots for 50 of 500 PCs. **D)** Outcomes from simulating 1000 parameter sets drawn from each of the two distributions. The black cross marks the value targeted by the NPE. See Table 3. In the baseline condition, the targeted outcome was well within support of the outcome distribution for all parameters. In the hyperexcitable condition, gamma and fast power targets were not within the interquartile range.

**Table 1:** The intrinsic parameters of the AdEx model populations. Square brackets indicate the lower and upper bounds of the uniform prior distribution. If a cell contains a single number, that parameter is held constant. Δ*T*, slope factor; *τ*_*w*_, adaptation time constant; *a*, subthreshold adaptation; *b*, spike-triggered adaptation; *V*_*r*_, spike triggered reset voltage; *C*, capacitance; *g*_*L*_, leak conductance; *E*_*L*_, resting membrane potential; *V*_*T*_, spike threshold. Note that *V*_*T*_ determines the point of exponential rise but is not the hard threshold for triggering the spike reset. A spike is triggered when the voltage is larger than 0 mV, see Equation 3.1; *n*, number of neurons in the population.

| Parameter | PC | AIN | NIN |
| --- | --- | --- | --- |
| $\Delta T$ | 0.8 mV | 5.5 mV | 3.0 mV |
| $\tau_w$ | 88 ms | 1 ms | 16 ms |
| $a$ | 0.8 nS | 2.0 nS | 1.8 nS |
| $b$ | 65 pA | 55 pA | 61 pA |
| $V_r$ | -53 mV | -54 mV | -54 mV |
| $C$ | [10 pF, 100 pF] | [96 pF, 209 pF] | [10 pF, 323 pF] |
| $g_L$ | [2.0 nS, 20.0 nS] | [2.0 nS, 6.4 nS] | [3.0 nS, 13.5 nS] |
| $E_L$ | [-80 mV, -64 mV] | [-65 mV, -51 mV] | [-70 mV, -58 mV] |
| $V_T$ | [-49 mV, -39 mV] | [-63 mV, -44 mV] | [-52 mV, -37 mV] |
| $n$ | 500 | [10, 100] | [10, 100] |

**Table 2:** The parameters of the Tsodyks-Markram synaptic connections. *A*_SE_, maximum synaptic conductance; *τ*_in_, time constant of synaptic decay; *τ*_rec_, synaptic resource recovery time constant; *U*_SE_, fraction of synaptic resources activated; *τ*_facil_, decay of facilitation time constant; *p* pairwise connection probability in %. NIN GJ NIN, is a gap junction connection, modeled as a fixed conductance between connected NINs.

| Connection | $A_{SE}$ | $\tau_{in}$ | $\tau_{rec}$ | $U_{SE}$ | $\tau_{facil}$ | $p$ |
| --- | --- | --- | --- | --- | --- | --- |
| Input → PC | 5.5 nS | 5.5 ms | 800 ms | 0.5 | 0 ms | 0.2 |
| Input → NIN | 7.0 nS | 6.3 ms | 800 ms | 0.5 | 0 ms | 0.1 |
| PC → PC | [0.0 nS, 50.0 nS] | 10.0 ms | 800 ms | 0.5 | 0 ms | [0.0, 0.3] |
| PC → AIN | [0.0 nS, 50.0 nS] | 4.0 ms | 130 ms | 0.03 | 530 ms | [0.0, 0.3] |
| PC → NIN | [0.0 nS, 50.0 nS] | 10 ms | 800 ms | 0.5 | 0 ms | [0.0, 0.3] |
| NIN → PC | [0.0 nS, 50.0 nS] | 5.5 ms | 800 ms | 0.5 | 0 ms | [0.0, 0.3] |
| NIN → NIN | [0.0 nS, 50.0 nS] | 2.5 ms | 800 ms | 0.5 | 0 ms | [0.0, 0.3] |
| AIN → PC | [0.0 nS, 50.0 nS] | 6.0 ms | 800 ms | 0.5 | 0 ms | [0.0, 0.3] |
| AIN → NIN | [0.0 nS, 50.0 nS] | 6.0 ms | 50 ms | 0.03 | 0 ms | [0.0, 0.3] |
| AIN → AIN | [0.0 nS, 50.0 nS] | 5.5 ms | 130 ms | 0.03 | 530 ms | [0.0, 0.3] |
| NIN GJ NIN | [0.0 nS, 4.0 nS] | - | - | - | - | [0.0, 0.3] |

We chose two network states as targets for simulation-based inference: A baseline (BL) condition that represents healthy activity and a hyperexcitable (HE) condition that represents pathological hyperactivity (Figure 1 C). To identify the parameter distributions underlying the BL and HE conditions, we trained a neural posterior estimator (NPE). Simulations with zero spikes were used by setting the ISI entropy, correlation and ISI CV to zero. We then used simulation-based calibration (Talts et al., 2018) to test the reliability of the posterior estimate *q*(*θ*|*x*) (*θ* are the parameters and *x* are the summary statistics,) and found that the posterior estimate was unreliable for several parameters (Figure Supp 1.1).

**Table 3:** The quantified outcomes of the simulator. ISI, interspike interval; CV, coefficient of variation.

| Outcome | Mean Rate<br>(Hz) | Mean ISI<br>Entropy | Theta<br>Power | Gamma<br>Power | Fast Power | Correlation | ISI CV |
| --- | --- | --- | --- | --- | --- | --- | --- |
| Baseline<br>(BL) | 18 | 2.7 | $1 \cdot 10^{-7}$ | $1 \cdot 10^{-8}$ | $1 \cdot 10^{-9}$ | 0.05 | 0.45 |
| Hyper-<br>excitable<br>(HE) | 75 | 1.0 | $9 \cdot 10^{-6}$ | $7 \cdot 10^{-9}$ | $6 \cdot 10^{-10}$ | 0.99 | 4.0 |

Because the NPE based solely on prior samples was unreliable, we used truncated sequential NPE (Deistler et al., 2022). This technique draws samples from posterior estimates to iteratively improve the estimate. The 2,620,000 prior samples were used to create the initial posterior estimate, and then 20,000 samples were drawn from the posterior estimates in 4 iterations. These simulations showed clearly distinct excitability levels of the simulator (Figure 1 C). Furthermore, the simulation outcomes from both posterior distributions were clearly distinct and covered most target outcomes (Figure 1 D). Only the gamma and fast power outcomes were not within the interquartile range of the HE condition.

To identify compensatory mechanisms that maintain the baseline excitability, we calculated the pairwise correlation coefficients between parameter samples from the posterior estimate. These correlations predict compensatory mechanisms, because they indicate that baseline activity can be maintained despite changes in one parameter if the other parameter changes accordingly. Since our model has 32 free parameters, 496 unique pairwise correlations exist. For each pair, we calculated the Spearman correlation coefficient from 200000 marginal posterior samples. Out of all pairs, only 19 had correlation coefficients larger than 0.1 or smaller than −0.1 (Figure 2 C).

**Figure 2:**
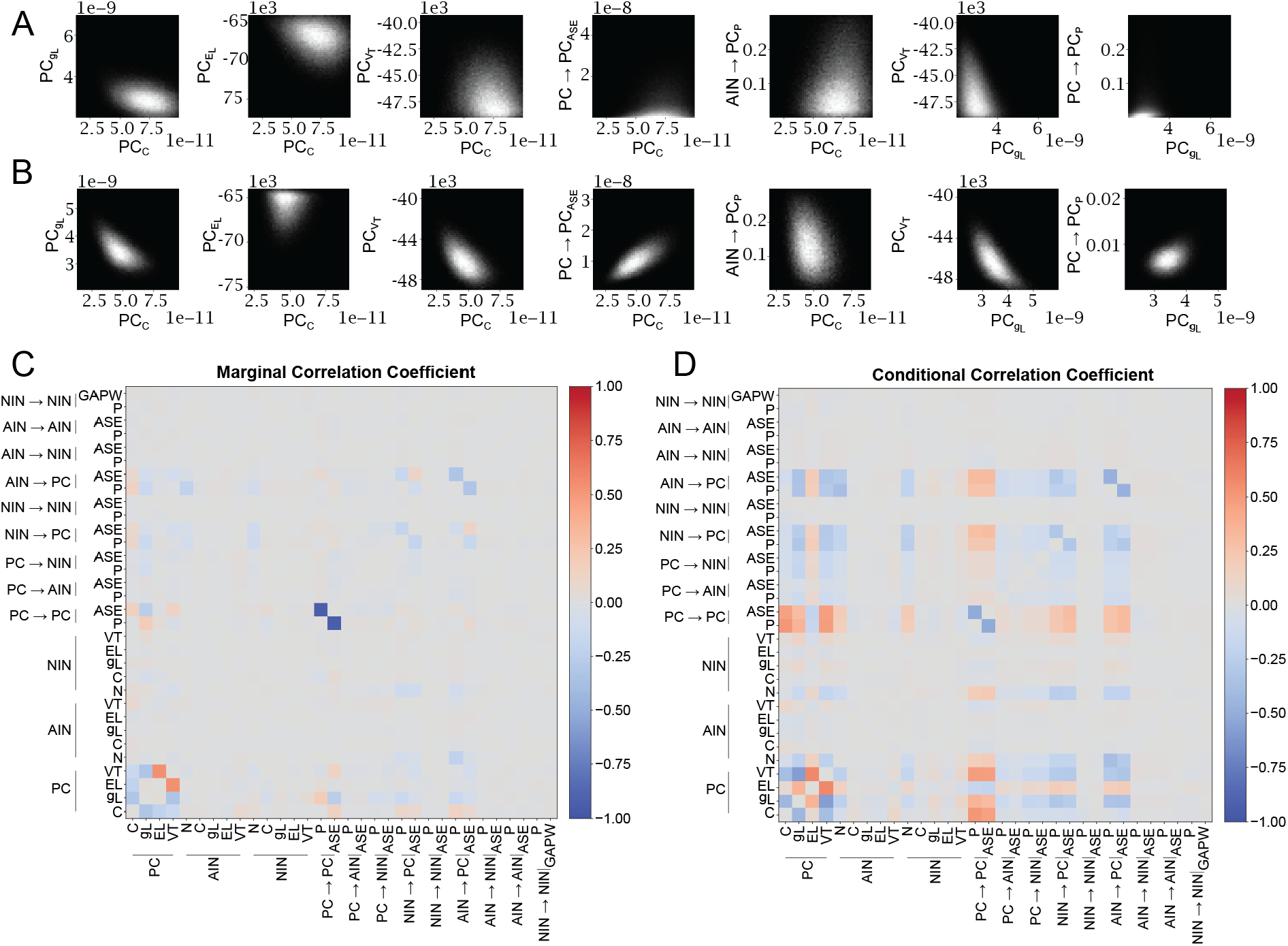
Various parameter pairs can compensate for each other to maintain baseline excitability. **A)** Shows seven representative 2D histograms of pairs with marginal correlation coefficients larger >0.1 or <−0.1. The maximum pixel values in order are: 813, 588, 525, 1718, 411, 863, 3328. All minimum values are zero. **B)** MAP conditional sample histograms for the same parameters as in A. The maximum pixel values in order are: 626, 606, 517, 627, 277, 519, 738. **C)** Shows the correlation coefficient of all parameter pairs as a diverging heatmap. The marginal correlation coefficients for the pairs shown in A from top left to bottom right, are: −0.31, −0.21, −0.13, 0.15, 0.13, −0.32, 0.19. **D)** Shows the correlation coefficients of samples drawn for each parameter pair while conditioning all other parameters on the map estimate. Correlation coefficients for the order in A from top left to bottom right, are: −0.43, 0.06, −0.41, 0.49, −0.01, −0.59, 0.42.

The largest negative correlation (−0.94) was between the PC connection probability (PC → PC_*p*_) and the connection strength (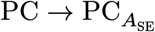). Both parameters were narrowly tuned and must be extremely small to maintain healthy dynamics (Figure 2 A). Probability and strength of connections from both interneuron types to PCs were also negatively correlated, albeit with smaller correlation coefficients (Figure 2 B), and they were not as narrowly constrained.

The largest positive correlation was between two intrinsic properties of PCs: the equilibrium potential of the leak current 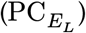 and the threshold (*V*_*T*_). The capacitance was negatively correlated with the other three intrinsic properties. Overall, this indicates that several putative compensatory mechanisms exist. But even correlated marginal samples are broadly distributed (Figure 2 A).

Marginal pairwise distributions are broad, because they are unconstrained by other parameters. But the posterior distribution can be used to constrain the other parameters. This led to narrower distributions (Figure 2 B) and, in most cases, larger absolute correlations (Figure 2 D). Furthermore, almost all parameters with meaningful marginal correlations also showed meaningful conditional correlations (Figure Supp 2.1). This shows that unconstrained and constrained correlations both suggest compensatory mechanisms.

Next, we set out to explore compensatory mechanisms for specific perturbations. By comparing two conditioned distributions, *p*(*θ*|*x*_BL_, *θ*_*i*_ = *k*) and *p*(*θ*|*x*_BL_, *θ*_*i*_ = *j*), we can estimate the compensatory mechanisms that maintain *x*_BL_ despite the condition. For example, if *θ*_*i*_ = *k* is a normal amount of interneurons and *θ*_*i*_ = *j* is the amount of interneurons expected in sclerotic tissue, we can find the parameters changes that can maintain healthy output despite the interneuron loss. We used this technique to investigate IN loss, recurrent excitation, and PC depolarization.

We sampled from the baseline posterior estimator with additional conditions on parameters representing known pathophysiological parameters (Figure 3). For interneuron loss, we conditioned the cell numbers in both interneuron populations (Normal: Equation 11 versus IN Loss: Equation 12). For recurrent excitation (called sprouting), we conditioned the connection probability PC → PC*p* (Normal: Equation 13 versus Sprouting: Equation 14). And for PC depolarization we conditioned 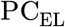 and 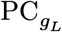 (Normal: Equation 15 versus Depolarized: Equation 16). While conditioning on these parameters, we sampled all other parameters and compared the normal with the pathophysiological samples. The difference between the distributions of a parameter estimates compensatory mechanisms because it shows how the parameter can change to maintain baseline activity despite the pathophysiology. We use the Kolmogorov-Smirnov (KS) test statistic to quantify the difference between distributions, where 0 indicates identical distributions and 1 indicates non-overlapping distributions.

**Figure 3:**
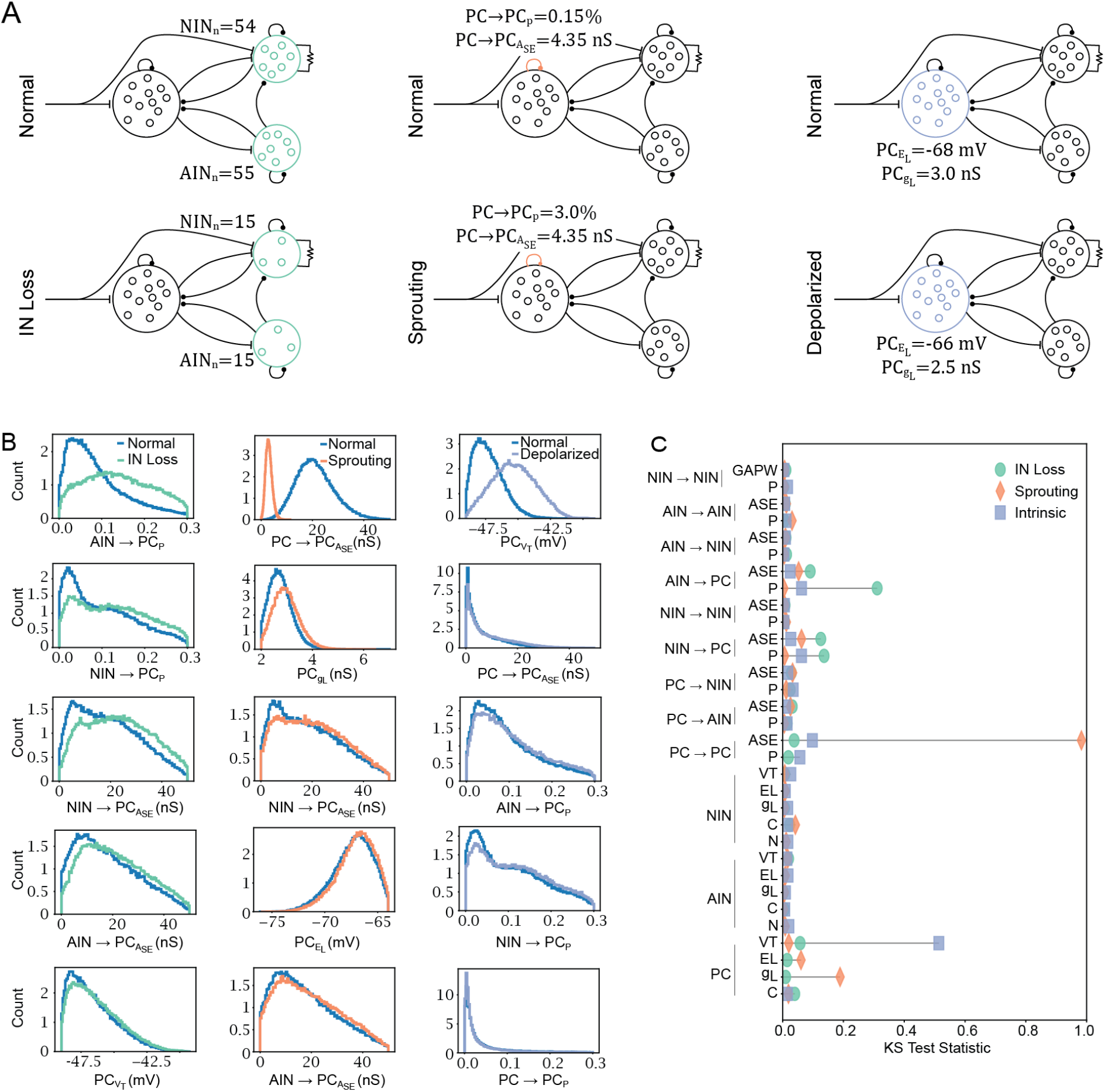
Putative compensatory parameters of specific pathophysiological conditions. **A)** A schematic illustration of the three conditions. IN loss compares 55 AINs and 54 NINs in each subpopulation (Normal, Equation 11) to 15 neurons in each population (IN Loss, Equation 12). Sprouting compares PC − PC_*p*_ of 0.15% (Normal, Equation 13) to 3% (Sprouting, Equation 14). Depolarized compares 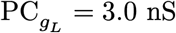, 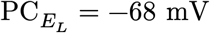 (Normal, Equation 15), with 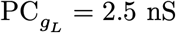, 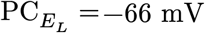 (Depolarized, Equation 16). **B)** shows the five parameters with the largest difference in each condition. KS test statistic for all parameters and conditions. **C)** Shows the KS statistics for all parameters and all conditions. KS test results for all parameters and conditions are in Table Supp 3.1.

We found that the different pathophysiological conditions had different compensatory profiles (Figure 3 A, B & C). For example, when the number of INs is constrained, increasing the AIN → PC_*p*_ connection probability maintains baseline excitability (Figure 3 A, leftmost). Figure 3 D shows the KS test statistic for each parameter in each condition. As expected from the pairwise correlations in Figure 2, the PC → PC connection strength has the largest KS statistic in the sprouting conditional. For IN Loss and Depolarization (Figure 3 A & C) on the other hand, the compensatory potential of the PC → PC connection is low. This shows that some compensatory parameters depend on other conditions. This is also true for IN loss and intrinsics. During IN loss, the connection probabilities from INs → PCs stand out as being particularly effective. In contrast, the threshold potential is uniquely positioned to maintain baseline excitability despite intrinsic depolarization.

The results presented so far came all from the same posterior density estimator. To assess reproducibility, we simulated 10,310,000 additional samples and trained two sequential NPEs. For amortized training 1,122,586 samples were randomly chosen for each and then the sequential loops were repeated as above. The replicates and the original data are overall highly correlated (Figure 4).

**Figure 4:**
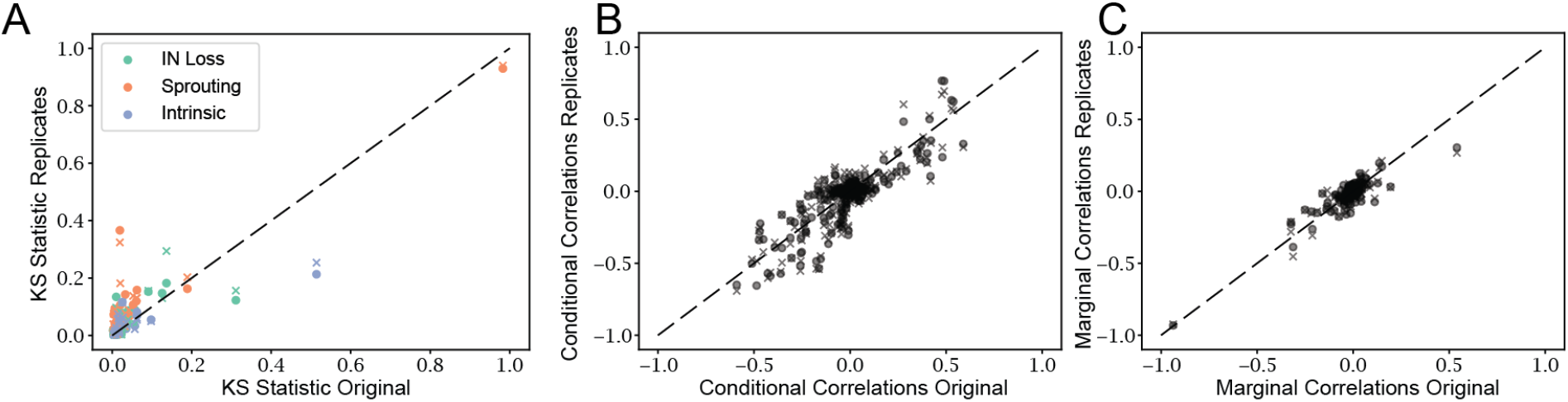
Replication of the posterior estimator with different training data. The dot and the x are results from separately trained estimators. **A)** Original KS statistics versus those from two separately trained replicates. See Figure 3 D for original data. **B)** Original conditional correlation coefficients versus those from the two replicates. See Figure 3 E for original data. **C)** Same as B but with marginal correlations. See Figure 3 F for original data. Correlation coefficients of A, B and C are 0.87, 0.84 and 0.89, respectively.

The above results focus on the mechanisms that maintain baseline excitability. Next, we investigated the putative compensatory mechanisms that can move the network from hyperexcitable to baseline (Figure 5 A). The parameters with the largest difference are across all three conditions the intrinsic and recurrent properties of PCs (Figure 5 B & C). The only two interneuron parameters that have compensatory potential are the capacitance of NINs and the threshold of AINs.

**Figure 5:**
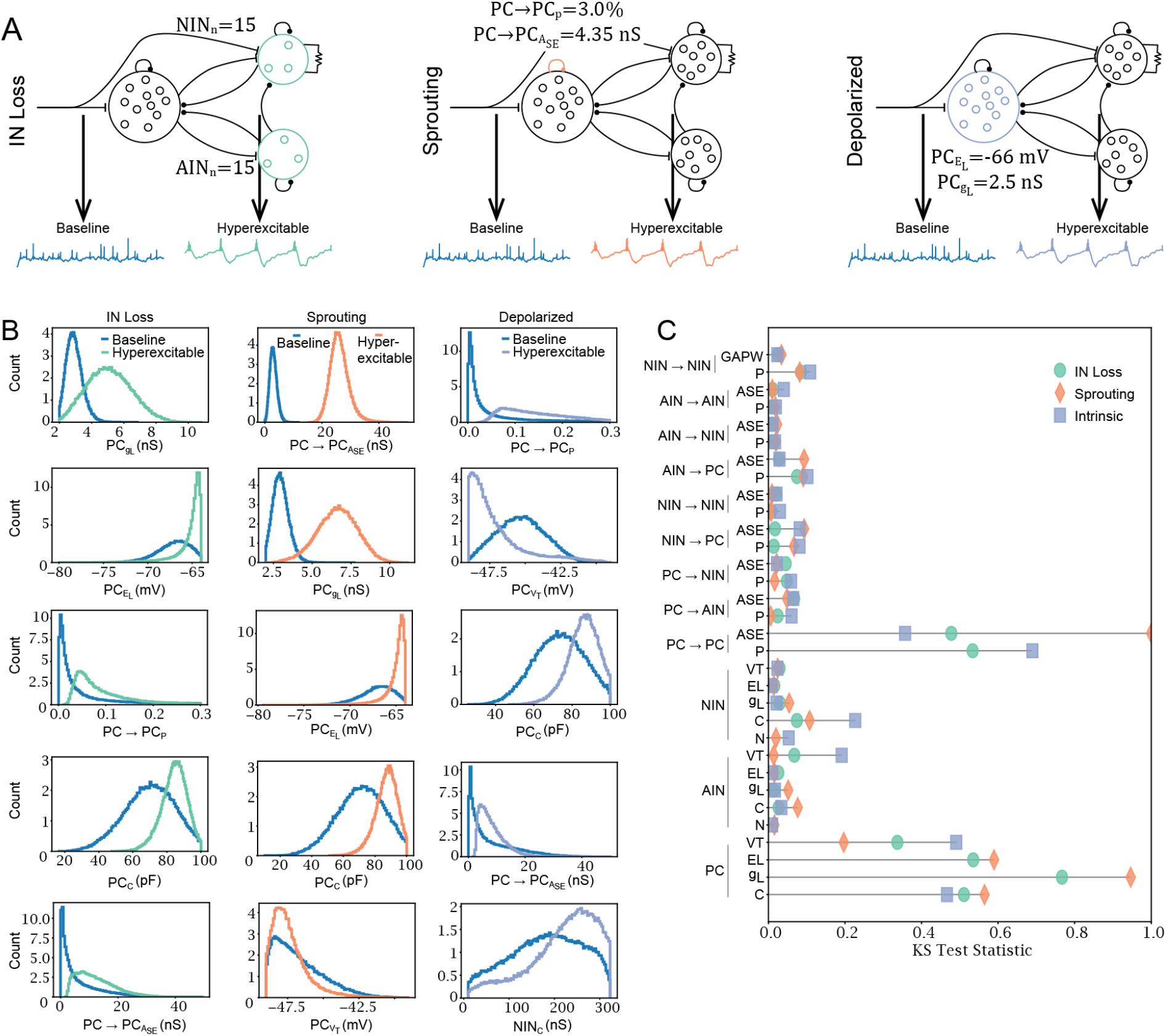
Putative compensatory parameters that can restore baseline activity given hyperexcitability. **A)** Schematic illustration of the three pathophysiological conditions and the comparison of baseline versus hyperexcitable network output. **B)** shows the five parameters with the largest test statistic. Each compares baseline and the hyperexcitable estimator for a specific pathophysiological condition. **C)** shows the KS test statistic calculated between the baseline and hyperexcitable posterior for each pathophysiological condition. KS test results for all parameters and conditions are in Table Supp 5.1.

We have used posterior density estimators to explore compensatory mechanisms. However, the main goal of a compensatory mechanism is to bring the system from the pathophysiological state back to the healthy state. In the simulator, we can directly test the effect of parameter changes on the output and, furthermore, whether the effect depends on the specific pathophysiological cause. To this end, we used the hyperexcitable posterior density estimator and drew samples given each pathophysiological condition: IN Loss, Sprouting and Depolarization.

To test the effect of a parameter on output, a single parameter was then systematically varied and simulated. If the pathophysiological condition influences the effect of a parameter on the output, we expect a change in the difference between conditions at some parameter point. The interaction of an ANOVA shows whether the interaction between the parameter change and any of the three pathophysiological conditions is significant. A statistically significant interaction indicates that the effect of a parameter on the outcome depends on the condition (or vice versa) and is thus specific to the pathophysiology. The results clearly show that the condition is important for the compensatory potential of many parameters (Figure 6). As expected, IN loss affects the influence of AIN → PC connectivity, and sprouting strongly affects 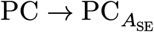. The effect of 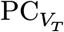 on a variety of outcomes is also strongly condition-dependent. Overall, this shows that the compensatory effect of many parameters depends on the specific pathophysiological condition.

**Figure 6:**
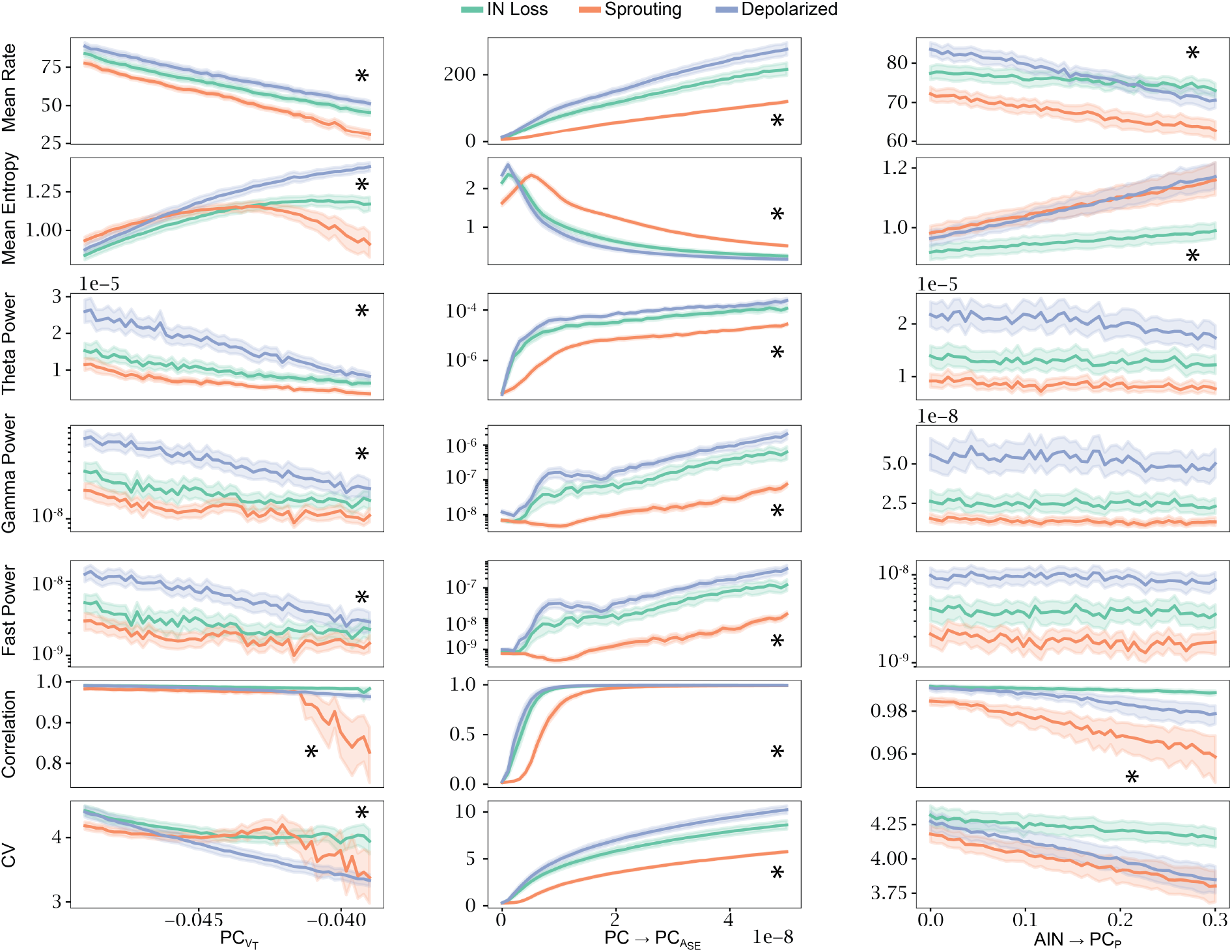
The pathophysiological condition changes the effect of parameters on simulated outcomes. The hyperexcitable posterior estimator was sampled with additional pathophysiological conditions. Then, each parameter was varied to cover 50 points in the parameters prior range and the parameters were simulated. Each of the 50 points on the x-axis contains 100 samples and the error bars show the 95% confidence interval. The asterisks indicate where the p-value of the interaction between parameter and condition is 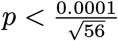. Figure Supp 6.1 shows the effect of more parameters and Table Supp 6.1 contains the p-values of the interactions.

## 3 Discussion

The strength of compensatory mechanisms depends on the underlying pathological causes. Using simulation-based inference, we quantify the ability of parameters to compensate for different causes of hyperexcitability. For example, we find that the threshold potential for exponential voltage rise (*V*_*T*_) is the best parameter to compensate for intrinsic depolarization. However, when compensating for interneuron loss or recurrent excitatory sprouting, *V*_*T*_ compensation is less effective. The exponential voltage rise in neurons is mediated by voltage-activated sodium channels, which are a crucial target of antiepileptic drugs (Kaplan et al., 2016). Our findings therefore predict that, in patients in whom seizures are not primarily caused by principal cell hyperexcitability, commonly used sodium channel blockers may be less effective. This could explain why sodium channel blockers do not control seizures in some patients.

Two other interesting parameters are the interneuron-to-principal cell connection probabilities (AIN → PC_*p*_ and NIN → PC_*p*_). Increasing the connection probability of either can compensate for intrinsic depolarization but also for the loss of interneurons. On one hand, fewer interneurons mean less overall inhibition and it makes sense that this can be compensated for by increasing connectivity. On the other hand, fewer interneurons means that the effect of the connectivity parameter should decrease, because the same increase in connection probability creates fewer overall synapses. We predict that even in sclerotic tissue with severe interneuron loss, increasing the number of inhibitory synapses has good compensatory potential. In vivo, thousands of interneurons remain even after interneuron loss in the pilocarpine or traumatic brain injury rat model of epilepsy (Huusko et al., 2015). The remaining interneurons could restore healthy activity by maintaining more synapses on principal cells. There are currently no epilepsy treatments that target inhibitory synapses directly, and increasing the number of inhibitory synapses without affecting excitatory synapses could prove difficult. A promising direction could be to use astrocyte-secreted factors such as neurocan, which specifically control inhibitory synaptogenesis (Irala et al., 2024).

While synaptogenesis has not yet been targeted for epilepsy treatments, interneuron transplantation to replace lost interneurons has been explored and proven effective in rodents (Hunt & Baraban, 2015). We find that the interneuron cell numbers, compared to synaptic and intrinsic parameters, do not compensate well for excitatory sprouting or intrinsic depolarization. Before using interneuron transplants, it could, therefore, be advisable to determine whether interneuron loss affects the epileptogenic region. This is especially important since some epileptogenic regions are non-sclerotic, having cell numbers similar to healthy controls (Blümcke et al., 2013). In these regions, seizures could be caused by other factors, and interneuron transplantation could be less effective.

Our results indicate that the pathophysiological mechanisms could be relevant for the effect of compensatory mechanisms, but measuring the number of interneurons, the number of recurrent excitatory synapses, or intrinsic excitability is currently impossible in living patients. It is, therefore, essential to develop methods to estimate these difficult-to-measure mechanistic features from measurable properties. Simulation-based inference can be used to estimate these mechanistic biomarkers. For example, Doorn et al. (2024) measured the activity of patient-derived neuronal cultures with multi-electrode arrays and used the posterior estimates of recurrent spiking neuronal network parameters conditioned on the activity as estimates of pathological mechanisms. This method shows some promise, but a gap remains between the activity in epileptic networks and the activity in cell cultures. Cell cultures are dominated by the details of the culturing technique and the patient’s genetics. This makes cell culture particularly useful for epilepsies with strong genetic components. In contrast, the activity of an epileptogenic network in vivo is more complex and depends on the patient’s genetics but also various protective or harmful environmental and developmental factors. Therefore, intracranial field recordings, as they are performed in pharmacoresistant epilepsies, could be necessary. Such recordings with phenomenological neuronal mass models, which simulate the average activity of coupled brain regions rather than the spiking of single neurons, have been used to estimate the epileptogenicity index (Makhalova et al., 2022). While this index does not describe a specific pathophysiological mechanism, it could be clinically meaningful. A clinical trial is ongoing to determine its efficacy in guiding surgical decisions on the resection of epileptogenic brain tissue (Jirsa et al., 2023). If neural mass models can provide meaningful estimates of epileptogenicity, spiking neuronal network models could provide estimates of pathophysiological mechanisms such as interneuron number.

Comparing conditional distributions has, to our knowledge, not been used to predict compensatory mechanisms. It is thus worth comparing it to the more common way of identifying compensation from the posterior correlations between parameters (Gonçalves et al., 2020). If we sample from the posterior *p*(*θ*|*x*), correlations between *θ*_*i*_ and *θ*_*j*_ indicate that if either of the parameters were changed, we could keep the probability of generating *x*-type output constant by changing the other parameter. This identifies broadly applicable compensatory mechanisms that work across the entire parameter range.

On the other hand, comparing *p*(*θ*_*i*_|*x, θ*_*j*_ = 5) to *p*(*θ*_*i*_|*x, θ*_*j*_ = 10) shows, how *θ*_*i*_ could change to maintain *x*-type dynamics, despite *θ*_*j*_ being perturbed from 5 to 10. The conditional also tests only the compensation of one parameter to another (the unconditioned compensates for the conditioned), whereas the correlations find compensation in both directions. Overall, correlations discover broadly applicable mechanisms, while conditionals identify specific mechanisms. Therefore, the conditional distributions could be more useful to generate hypotheses about the underlying mechanisms than the unspecific correlations.

In summary, we show that the pathophysiological mechanisms underlying hyperexcitability change the efficacy of compensatory mechanisms to restore normal excitability in a spiking neuronal network model. These findings suggest that these mechanisms and a biologically detailed simulator could be used to generate hypotheses about the effects of compensatory mechanisms, such as voltage-gated sodium channel blockers or interneuron transplants. Simulation-based exploration of possible treatment outcomes would be a major advance for the study of brain disorders.

## 4 Materials and Methods

### 4.1 The Spiking Neuronal Network Simulator

To explore the compensatory mechanisms of hyperexcitability with simulation-based inference, we implemented a phenomenological spiking neuronal network simulator (Figure 7). We used adaptive exponential leaky integrate-and-fire neurons (AdEx) (Brette & Gerstner, 2005) to model one excitatory and two inhibitory populations connected with conductance-based Tsodyks-Markram synapses to model short-term synaptic plasticity (Tsodyks et al., 1998). The voltage of the AdEx neuron changes according to Equation 1:

**Figure 7:**
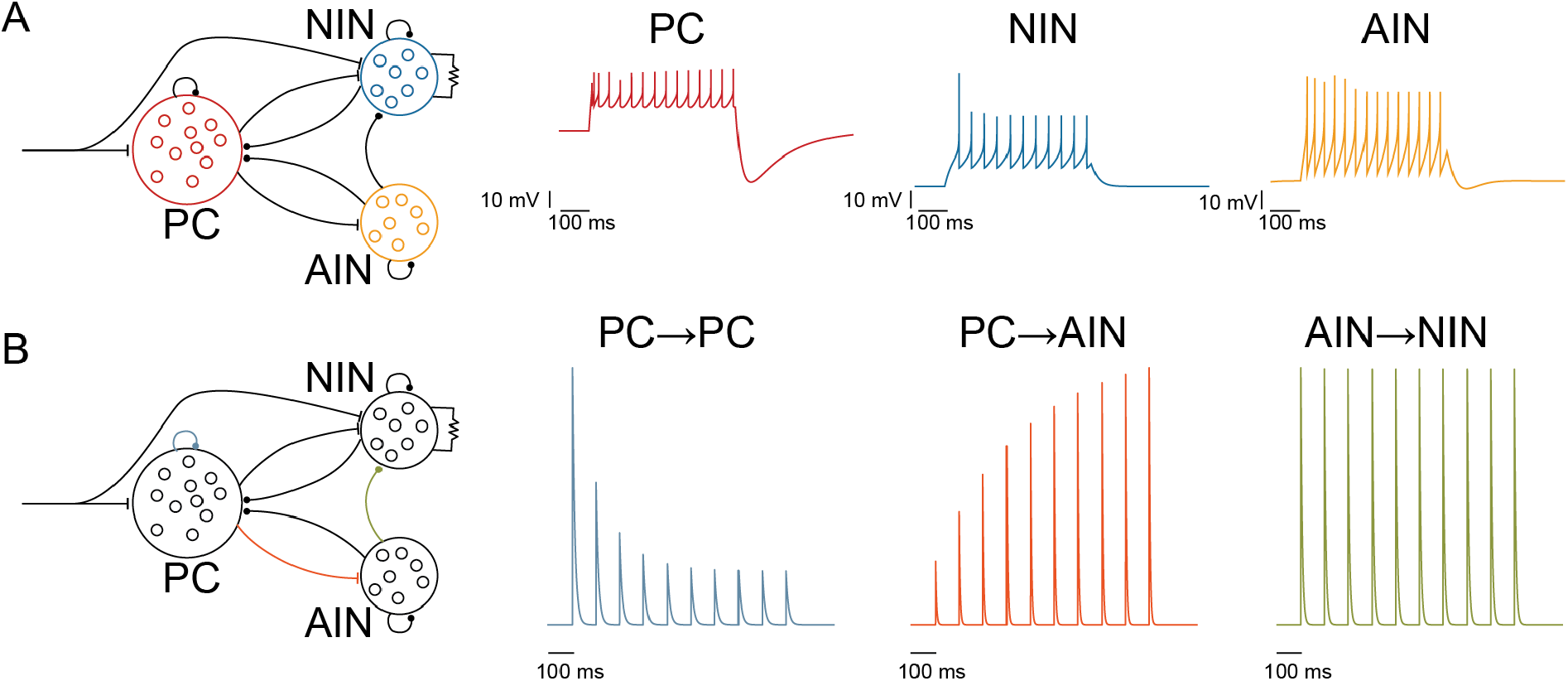
Representative illustrations of the AdEx neuron model (**A**) and the Tsodyks-Markram synapses (**B**) properties of the microcircuit simulator. In this example, parameters were fixed to the MAP of the baseline condition.

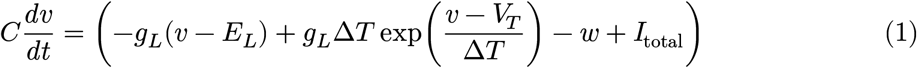

*C* is the membrane capacitance, *v* is the membrane voltage, *t* it the current time, *g*_*L*_ is the leak conductance, *E*_*L*_ is the leak conductance’s equilibrium potential, Δ*T* is the slope factor of the exponential term, *V*_*T*_ is the exponential threshold, *I*_total_ is the summed synaptic input (Equation 4), and *w* is the dynamic adaptation current:

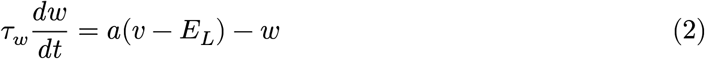

*τ*_*w*_ is the adaptation time constant and *a* is the magnitude of the subthreshold adaptation. When the membrane voltage is larger than 0, *v* and *w* are reset according to Equation 3.1 and Equation 3.2, respectively:

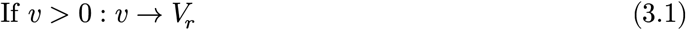

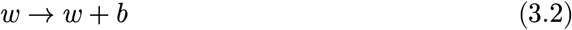

*V*_*r*_ is the reset voltage and *b* is the spike-triggered adaptation. All neuron model constants and parameters are in Table 1.

A neuron received its input through *I*_total_:

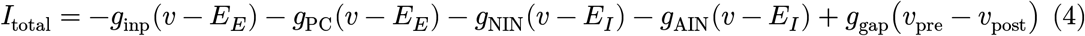

*g*_PC_, *g*_NIN &_ *g*_AIN_ are the conductances from the presynaptic principal cells (PC), non-adapting interneurons (NIN) and adapting interneurons (AIN), respectively. *g*_INP_ is the excitatory conductance from a homogeneous Poisson spike train. *g*_GAP_ is a constant conductance that exists only between some NINs to simulate symmetric gap junction connections.

The conductances and the synaptic resources changed as in Equation 5.3, Equation 5.2, Equation 5.5, Equation 5.4 and Equation 5.1 (Tsodyks et al., 1998).

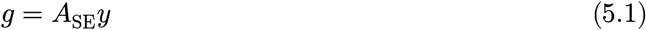

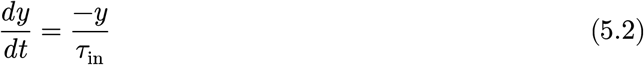

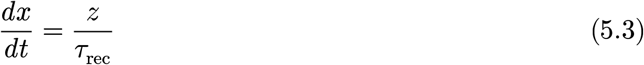

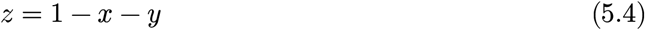

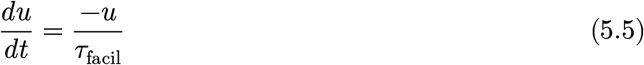

*x, y*, and *z* are fractions of synaptic resources. Resources in the recovered state (*x*) transition to the active state (*y*) when a presynaptic spike occurs (Equation 6.2). Resources in the active state (*y*) contribute to the synaptic conductance scaled by the synaptic weight *A*_SE_ (Equation 5.1) and transition to the inactive state (*z*) with time constant *τ*_in_ (Equation 5.2). Resources transition to the recovered state (*x*) from the inactive state (*z*) with the time constant *τ*_rec_ (Equation 5.3). Since the synaptic resources are fractions, the inactive resources can be calculate given the other two resources (Equation 5.4). The facilitation value *u* decays with time constant *τ*_facil_, and it facilitates because it increases when a spike occurs (Equation 6.1), and it scales the transition to the active fraction (Equation 6.2 & Equation 6.3). The parameter *U*_SE_ determines the strength of the facilitation. If *τ*_facil_ was set to 0, Equation 5.5 was deleted from the model, and *u* was kept constant at *U*_SE_. Table 2 shows the synaptic parameters for all connections.

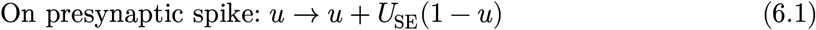

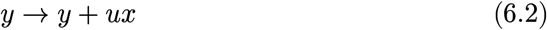

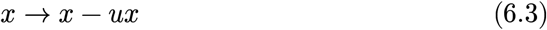

The model consists of three different populations: principal cells (PCs), adapting interneurons (AINs), and non-adapting interneurons (NINs) (Figure 7 A). Their parameters are largely taken from Naud et al. (2008). The AdEx model has nine parameters, and we additionally varied the number of neurons in the interneuron populations (see Table 1 for all parameters). We kept Δ*T, τ*_*w*_, *a, b*, and *V*_*r*_ constant to keep the parameter space tractable for simulation-based inference (SBI) and to keep qualitative changes of firing patterns to a minimum. That left 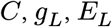and *V*_*T*_ subject to SBI. This totals twelve free neuronal parameters plus the cell numbers of the two interneuron populations.

The neuronal populations and their homogeneous Poisson input were connected through 10 types of synapses and 1 gap junction (Figure 7 B). The homogeneous Poisson input innervated the PCs and the NINs. All parameters of both inputs were fixed. For the other chemical synapses we fixed the parameters *τ*_in_, *τ*_rec_, *U*_SE_ and *τ*_facil_. The connection probability (in %) and the strength were varied during SBI. The chemical connections in the network were PC → PC, PC → aIN, PC → NIN, NIN → PC, NIN → NIN, AIN → PC, AIN → NIN, and AIN → AIN. Gap junctions connected NINs to other NINs. Therefore, eighteen connectivity parameters were varied during SBI.

We implemented the model with Brian2 (Stimberg et al., 2019). All simulations ran for 1 s with a temporal resolution of 0.1 ms.

### 4.2 Summary Statistics

We performed inference given seven summary statistics, all of which were calculated on the PCs (Table 3). The mean rate was calculated from all PCs, including silent ones. To calculate the mean interspike interval (ISI) entropy, we calculated the histogram of the ISIs in bins of 1 ms. Theta, gamma and fast power were calculated from the power spectral density (PSD). Theta power was calculated by summing the PSD between 8 Hz and 12 Hz, Gamma power between 30 Hz and 100 Hz and fast power between 100 and 150. The correlation was calculated between the convolved spike trains of 50 randomly chosen PCs and then averaged. For the average pairwise Pearson correlation, spike trains were convolved with as Gaussian kernel of 10 ms width. Pairs where one of the neurons had zero spikes were excluded from the average. The ISI coefficient of variance (CV) for each cell was calculated by dividing the mean of the ISI by the standard deviation of the ISI. Then, CVs of all cells were averaged, excluding cells with undefined CVs (less than two spikes or zero standard deviation).

### 4.3 Simulation-Based Inference and Quality Checks

For SBI we used neural posterior estimation (NPE; Gonçalves et al. (2020); Papamakarios & Murray (2018)) with the SNPE_C inference implementation from the sbi Python package 0.22.0 (Tejero-Cantero et al., 2020). Initially, 2620000 parameter samples were drawn from the prior, then simulated, and the parameter-outcome pairs were used to train a neuronal density estimator with default parameters. For sequential inference, we used truncated sequential NPE (Deistler et al., 2022). The posterior estimate was truncated at the 0.0001 quantile in four sequential rounds, with 20000 simulations in each round. 1000000 samples were used to estimate the support. Undefined summary statistics (ISI Entropy, correlation or ISI CV) were set to 0. To calculate marginal pairwise Spearman correlations, 200000 samples were drawn from the baseline posterior distribution. To calculate the conditional correlation coefficients we conditioned other parameters on the maximum-a-posteriori (MAP) estimate.

To assess the reproducibility of the posterior estimation (Figure 4), two estimators were trained. The hyperparameters of the training were as above, but to train the estimators, 1,122,586 simulations were randomly drawn from a pool of 10,310,000 that included the samples used to train the original estimator. Training and inference of these estimators was performed with the 0.24.0 version of the sbi Python package.

### 4.4 Specific Conditional Baseline Distributions

We introduced specific conditions on the posterior distribution to investigate the mechanisms that can compensate for interneuron loss, excitatory synaptic sprouting, and intrinsic hyperexcitability. To investigate interneuron loss we conditioned the number of AINs (AIN_*n*_) and the number of NINs (NIN_*n*_):

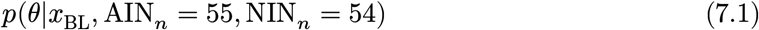

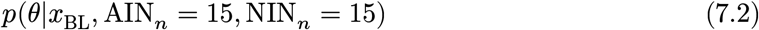

Equation 7.2 produces healthy dynamics (*x*_BL_), despite interneuron loss and Equation 7.1 is the healthy control, which produces healthy dynamics with a normal number of interneurons. *θ* are the unconditioned parameters. 55 and 54 are the MAP estimates for the interneuron counts of the marginal baseline posterior.

To investigate excitatory synaptic sprouting we conditioned the PC to PC connection probability (PC → PC_*p*_) and strength 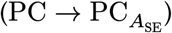 :

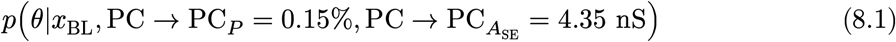

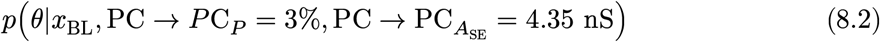

Equation 8.2 produces healthy dynamics despite excitatory sprouting. The connection probability 3% is double the baseline MAP estimate of 1.5%. For comparison we chose an order of magnitude smaller percentage points for Equation 8.1. The connection strength 4.35 nS is the MAP estimate of the marginal BL posterior.

To investigate intrinsic changes in principal cell excitability, we conditioned the PC leak conductance 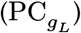 and PC equilibrium potential 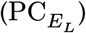 :

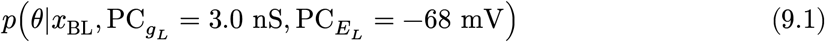

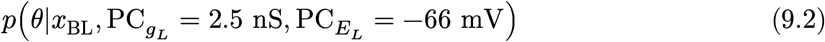

These represent a minor increase of input resistance and resting membrane potential in principal cells. MAP estimate for baseline output of the two parameters was 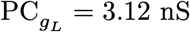 and 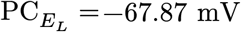.

To compare conditional distributions of parameters, 100000 samples were drawn and we used the Kolmogorov-Smirnov test to compare the healthy to the perturbed conditional.

### 4.5 Conditional Baseline versus Hyperexcitable Distributions

To compare the parameters that produce baseline (*x*_BL_) and hyperexcitable (*x*_HE_) activity despite pathological perturbations (Figure 5), we used MCMC sampling to draw 100000 samples and compared the distributions with the Kolmogorov-Smirnov test.

To predict compensatory mechanisms of interneuron loss, we conditioned AINs (AIN_*n*_) and the number of NINs (NIN_*n*_) in the baseline (Equation 10.1) and hyperexcitable condition (Equation 10.2):

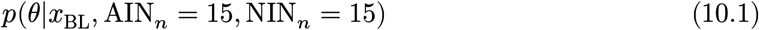

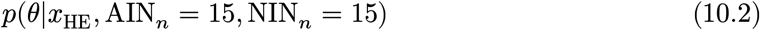

For synaptic sprouting, we conditioned the PC to PC connection probability 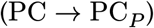 and strength 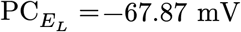in the baseline (Equation 11.1) and hyperexcitable (Equation 11.2) condition:

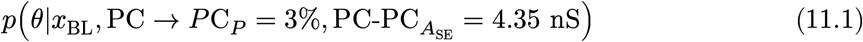

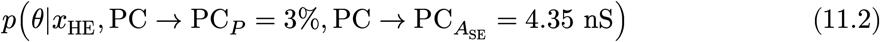

For depolarization, we conditioned the PC leak conductance 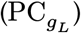 and PC equilibrium potential 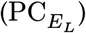 in the baseline (Equation 12.1) and hyperexcitable (Equation 12.2) condition:

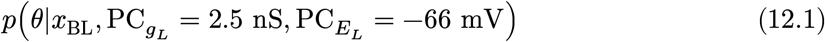

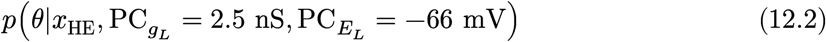

### 4.6 Compensatory Effects on Simulator Output

Finally, we sampled parameters from the hyperexcitable condition given pathological perturbations (see Equation 10.2, Equation 11.2 & Equation 12.2) and ran simulations to determine the effects of parameters on the outcome variables (Figure 6). In each condition, we drew 100 samples. For each parameter and each sample, we varied the parameter linearly to obtain 50 values spanning the lowest to the highest prior value.

### 4.7 Software

Prior samples were drawn and simulated on the Okinawa Institute of Science High Performance computing resources running CentOS 8. with the 4. version of the Linux kernel. The brian2 version was 2.5.0.3 (Stimberg et al., 2019). The Python version was 3.9.18.

The neural density estimator training, sequential NPE and conditional sampling were run on a Windows 10 computer with brian2 2.7.1. The Python version was 3.10.14.

Plotting was done with matplotlib 3.9.1 (Hunter, 2007) or seaborn 0.13.2 (Waskom, 2021). pandas 2.2.3 was used for data wrangling (McKinney, 2010). statsmodels 0.14.4 was used for ANOVA test (Seabold & Perktold, 2010). scipy 1.13.1 was used for Kolmogorov-Smirnov tests and Spearman correlation coefficient (Virtanen et al., 2020).

## 5 Code and Data Availability

The Python code to run the simulator, train the NPE, infer posteriors, analyze the data and plot the figures is available on GitHub (https://github.com/danielmk/hyperexcitability_sbi). The simulated data and the trained NPEs are publicly available at https://doi.org/10.5281/zenodo.20152410.

## 6 Acknowledgements

We are grateful for the help and support provided by the Scientific Computing and Data Analysis section of Core Facilities at OIST. We thank Milena Menezes Carvalho, Joanna Komorowska-Müller and Gastón Sivori for helpful comments on the manuscript. This work was supported by JSPS KAKENHI grant no. JP23H05476 to T.F.

## Notes

### Competing Interest Statement

The authors have declared no competing interest.

### Summary of Updates

Changed parts of the text. Fixed typos. Added a supplemental figure that shows the simulation-based calibration results of the amortized estimator.

