## supplemental_figures for "Compensation of Hyperexcitability with Simulation-Based Inference"

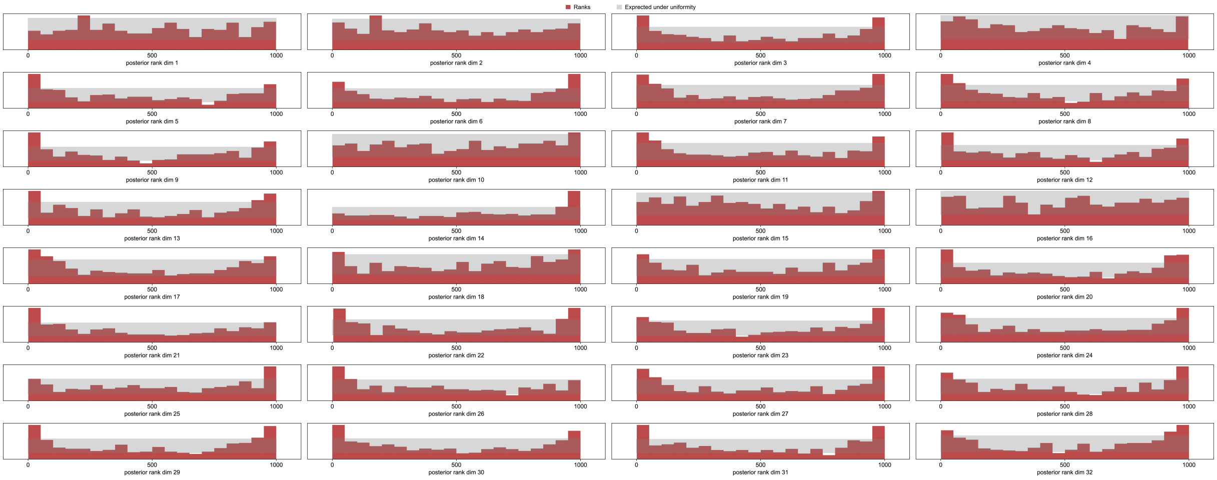


Supp 1.1 Simulation-based calibration results of an amortized posterior estimator. The red bars indicate the ranks distribution of ranks. Each histogram is for a specific parameter. If the posterior estimates were well calibrated, the rank distributions would be uniformly distributed. The grey background indicates the 99% confidence interval of a uniform distribution given the number of samples. The p-values of the null-hypothesis that the ranks are drawn from a uniform distribution were as follows: 0.6784; 0.1548; 0.0008; 0.0245; 0.0000; 0.0000; 0.0001; 0.0000; 0.0000; 0.0801; 0.0000; 0.0023; 0.0001; 0.0000; 0.0296; 0.2425; 0.0000; 0.0034; 0.0010; 0.0000; 0.0000; 0.0004; 0.0003; 0.0007; 0.0015; 0.0001; 0.0000; 0.0008; 0.0000; 0.0000; 0.0000; 0.0000. Since several of the p-values were well below 0.001, we concluded the amortized estimate was mis-calibrated.

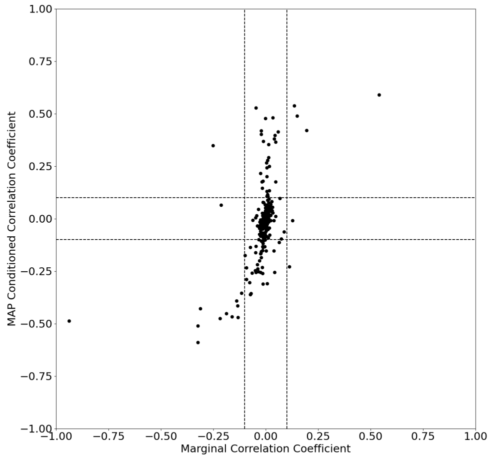


Supp 2.1 Marginal versus MAP conditioned correlation coefficient. Only $PC_C$ vs $PC_{E_L}$ was not meaningfully correlated with marginal sampling but became non-meaningfully correlated with conditioned sampling. Interestingly $PC_{g_L}$ vs PC-$PC_{A_{SE}}$ switched from negative correlation with marginal sampling to positive correlation with conditional sampling. The NIN - $PC_{A_{SE}}$ vs AIN - $PC_{A_{SE}}$ switched from positive in the marginal to negative in the conditional.

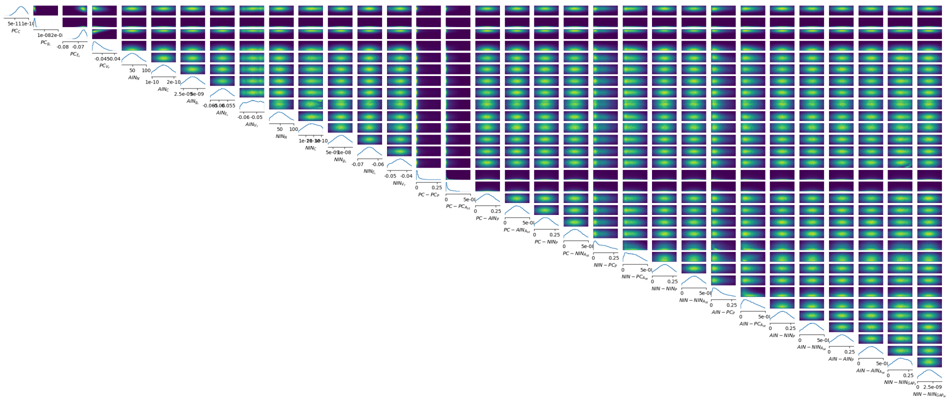

Supp 2.2 Full baseline marginal posterior.

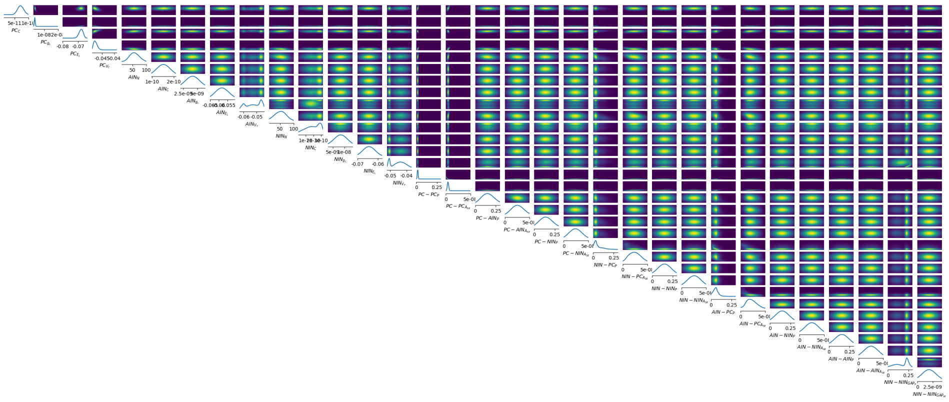

Supp 2.3 Full baseline MAP conditioned posterior.

Supp 3.1 Outcomes of the Kolmogorov-Smirnov test for the different conditionals of the baseline posterior.

| | IN Loss Statistic | IN Loss p Value | Instrinsic Statistic | Intrinsic p Value | Sprouting Statistic | Sprouting p Value |
|---|---|---|---|---|---|---|
| PC_C | 3.99E-02 | 1.05E-69 | 1.58E-02 | 2.51E-11 | 1.92E-02 | 2.24E-16 |
| PC_{g_L} | 9.97E-03 | 9.58E-05 | | | 1.89E-01 | 0.00E+00 |
| PC_{E_L} | 1.51E-02 | 2.19E-10 | | | 6.04E-02 | 4.10E-159 |
| PC_{V_T} | 5.75E-02 | 6.18E-144 | 5.14E-01 | 0.00E+00 | 2.03E-02 | 3.05E-18 |
| AIN_N | | | 2.06E-02 | 8.98E-19 | 8.54E-03 | 1.35E-03 |
| AIN_C | 5.69E-03 | 7.82E-02 | 6.84E-03 | 1.85E-02 | 2.80E-03 | 8.27E-01 |
| AIN_{g_L} | 3.21E-03 | 6.80E-01 | 9.48E-03 | 2.48E-04 | 5.90E-03 | 6.13E-02 |
| AIN_{E_L} | 2.31E-03 | 9.52E-01 | 1.79E-02 | 2.15E-14 | 8.98E-03 | 6.25E-04 |
| AIN_{V_T} | 1.97E-02 | 2.64E-17 | 1.45E-02 | 1.38E-09 | 1.67E-02 | 1.80E-12 |
| NIN_N | | | 1.73E-02 | 1.79E-13 | 1.17E-02 | 2.25E-06 |
| NIN_C | 2.32E-02 | 1.04E-23 | 1.70E-02 | 5.18E-13 | 4.19E-02 | 6.79E-77 |
| NIN_{g_L} | 6.43E-03 | 3.19E-02 | 1.68E-02 | 1.09E-12 | 5.61E-03 | 8.56E-02 |
| NIN_{E_L} | 4.53E-03 | 2.56E-01 | 8.98E-03 | 6.25E-04 | 8.80E-03 | 8.62E-04 |
| NIN_{V_T} | 7.22E-03 | 1.08E-02 | 2.70E-02 | 4.02E-32 | 7.06E-03 | 1.36E-02 |
| PC-PC_P | 1.89E-02 | 5.00E-16 | 5.68E-02 | 1.85E-140 | | |
| PC-PC_{A_{SE}} | 3.78E-02 | 1.31E-62 | 9.77E-02 | 0.00E+00 | 9.83E-01 | 0.00E+00 |
| PC-AIN_P | 1.52E-02 | 1.67E-10 | 1.49E-02 | 4.25E-10 | 5.51E-03 | 9.57E-02 |
| PC-AIN_{A_{SE}} | 3.17E-02 | 4.96E-44 | 1.01E-02 | 8.16E-05 | 2.59E-02 | 1.43E-29 |
| PC-NIN_P | 2.52E-02 | 6.61E-28 | 3.48E-02 | 4.49E-53 | 1.16E-02 | 2.65E-06 |
| PC-NIN_{A_{SE}} | 2.87E-02 | 3.88E-36 | 1.69E-02 | 7.53E-13 | 3.27E-02 | 5.71E-47 |
| NIN-PC_P | 1.36E-01 | 0.00E+00 | 6.20E-02 | 1.79E-167 | 6.32E-03 | 3.67E-02 |
| NIN-PC_{A_{SE}} | 1.26E-01 | 0.00E+00 | 2.68E-02 | 1.12E-31 | 6.21E-02 | 3.55E-168 |
| NIN-NIN_P | 7.16E-03 | 1.18E-02 | 4.49E-03 | 2.65E-01 | 1.22E-02 | 7.33E-07 |
| NIN-NIN_{A_{SE}} | 8.07E-03 | 2.95E-03 | 6.25E-03 | 4.01E-02 | 6.74E-03 | 2.12E-02 |
| AIN-PC_P | 3.11E-01 | 0.00E+00 | 6.20E-02 | 9.60E-168 | 3.85E-03 | 4.48E-01 |
| AIN-PC_{A_{SE}} | 9.08E-02 | 0.00E+00 | 2.53E-02 | 2.66E-28 | 5.26E-02 | 1.02E-120 |
| AIN-NIN_P | 1.31E-02 | 6.63E-08 | 4.35E-03 | 3.00E-01 | 5.26E-03 | 1.25E-01 |
| AIN-NIN_{A_{SE}} | 1.13E-02 | 5.91E-06 | 9.54E-03 | 2.22E-04 | 9.29E-03 | 3.55E-04 |
| AIN-AIN_P | 5.59E-03 | 8.76E-02 | 1.15E-02 | 3.34E-06 | 3.09E-02 | 5.12E-42 |
| AIN-AIN_{A_{SE}} | 9.07E-03 | 5.32E-04 | 7.14E-03 | 1.22E-02 | 1.24E-02 | 4.16E-07 |
| NIN-NIN_{GAP_P} | 4.06E-03 | 3.81E-01 | 1.63E-02 | 5.18E-12 | 5.79E-03 | 6.97E-02 |
| NIN-NIN_{GAP_W} | 1.11E-02 | 8.85E-06 | 4.56E-03 | 2.49E-01 | 6.66E-03 | 2.36E-02 |

Table Supp 5.1: Outcomes of the Kolmogorov-Smirnov test for comparison of the baseline and the hyperexcitable condition.

| | IN Loss Statistic | IN Loss p Value | Instrinsic Statistic | Intrinsic p Value | Sprouting Statistic | Sprouting p Value |
|---|---|---|---|---|---|---|
| PC_C | 5.11E-01 | 0.00E+00 | 4.67E-01 | 0.00E+00 | 5.65E-01 | 0.00E+00 |
| PC_{g_L} | 7.67E-01 | 0.00E+00 | | | 9.46E-01 | 0.00E+00 |
| PC_{E_L} | 5.35E-01 | 0.00E+00 | | | 5.89E-01 | 0.00E+00 |
| PC_{V_T} | 3.37E-01 | 0.00E+00 | 4.91E-01 | 0.00E+00 | 1.97E-01 | 0.00E+00 |
| AIN_N | | | 9.53E-03 | 2.26E-04 | 1.56E-02 | 5.01E-11 |
| AIN_C | 2.66E-02 | 3.09E-31 | 3.38E-02 | 4.93E-50 | 7.65E-02 | 7.16E-255 |
| AIN_{g_L} | 1.42E-02 | 3.56E-09 | 1.71E-02 | 3.56E-13 | 5.17E-02 | 1.11E-116 |
| AIN_{E_L} | 2.59E-02 | 1.76E-29 | 1.01E-02 | 7.68E-05 | 1.54E-02 | 9.04E-11 |
| AIN_{V_T} | 6.76E-02 | 4.75E-199 | 1.91E-01 | 0.00E+00 | 1.37E-02 | 1.52E-08 |
| NIN_N | | | 5.31E-02 | 6.33E-123 | 1.97E-02 | 2.86E-17 |
| NIN_C | 7.44E-02 | 3.32E-241 | 2.27E-01 | 0.00E+00 | 1.08E-01 | 0.00E+00 |
| NIN_{g_L} | 2.95E-02 | 2.44E-38 | 2.11E-02 | 8.70E-20 | 5.42E-02 | 4.19E-128 |
| NIN_{E_L} | 1.55E-02 | 7.28E-11 | 8.38E-03 | 1.77E-03 | 1.40E-02 | 5.76E-09 |
| NIN_{V_T} | 2.91E-02 | 2.56E-37 | 2.35E-02 | 1.76E-24 | 2.42E-02 | 7.92E-26 |
| PC-PC_P | 5.33E-01 | 0.00E+00 | 6.90E-01 | 0.00E+00 | | |
| PC-PC_{A_{SE}} | 4.78E-01 | 0.00E+00 | 3.57E-01 | 0.00E+00 | 1.00E+00 | 0.00E+00 |
| PC-AIN_P | 2.41E-02 | 1.06E-25 | 6.04E-02 | 4.10E-159 | 5.67E-03 | 8.00E-02 |
| PC-AIN_{A_{SE}} | 6.73E-02 | 2.74E-197 | 6.47E-02 | 1.23E-182 | 4.85E-02 | 8.92E-103 |
| PC-NIN_P | 4.82E-02 | 2.40E-101 | 5.91E-02 | 2.36E-152 | 1.63E-02 | 6.30E-12 |
| PC-NIN_{A_{SE}} | 4.51E-02 | 5.20E-89 | 2.21E-02 | 1.44E-21 | 2.15E-02 | 1.72E-20 |
| NIN-PC_P | 1.39E-02 | 8.27E-09 | 8.15E-02 | 1.74E-289 | 6.68E-02 | 1.98E-194 |
| NIN-PC_{A_{SE}} | 1.73E-02 | 2.05E-13 | 8.18E-02 | 2.90E-291 | 9.33E-02 | 0.00E+00 |
| NIN-NIN_P | 7.65E-03 | 5.72E-03 | 3.01E-02 | 6.80E-40 | 6.64E-03 | 2.42E-02 |
| NIN-NIN_{A_{SE}} | 2.06E-02 | 7.61E-19 | 2.09E-02 | 1.78E-19 | 9.20E-03 | 4.19E-04 |
| AIN-PC_P | 7.47E-02 | 5.87E-243 | 1.02E-01 | 0.00E+00 | 9.10E-02 | 0.00E+00 |
| AIN-PC_{A_{SE}} | 2.79E-02 | 3.39E-34 | 2.89E-02 | 8.18E-37 | 9.33E-02 | 0.00E+00 |
| AIN-NIN_P | 1.15E-02 | 3.84E-06 | 1.72E-02 | 2.36E-13 | 1.96E-02 | 3.92E-17 |
| AIN-NIN_{A_{SE}} | 8.71E-03 | 1.01E-03 | 1.07E-02 | 2.12E-05 | 2.28E-02 | 5.96E-23 |
| AIN-AIN_P | 8.71E-03 | 1.01E-03 | 1.95E-02 | 6.77E-17 | 1.84E-02 | 3.76E-15 |
| AIN-AIN_{A_{SE}} | 1.11E-02 | 9.67E-06 | 4.03E-02 | 5.83E-71 | 1.01E-02 | 7.53E-05 |
| NIN-NIN_{GAP_P} | 8.48E-02 | 4.19E-313 | 1.09E-01 | 0.00E+00 | 8.18E-02 | 2.09E-291 |
| NIN-NIN_{GAP_W} | 2.48E-02 | 3.27E-27 | 2.33E-02 | 5.95E-24 | 3.40E-02 | 1.67E-50 |

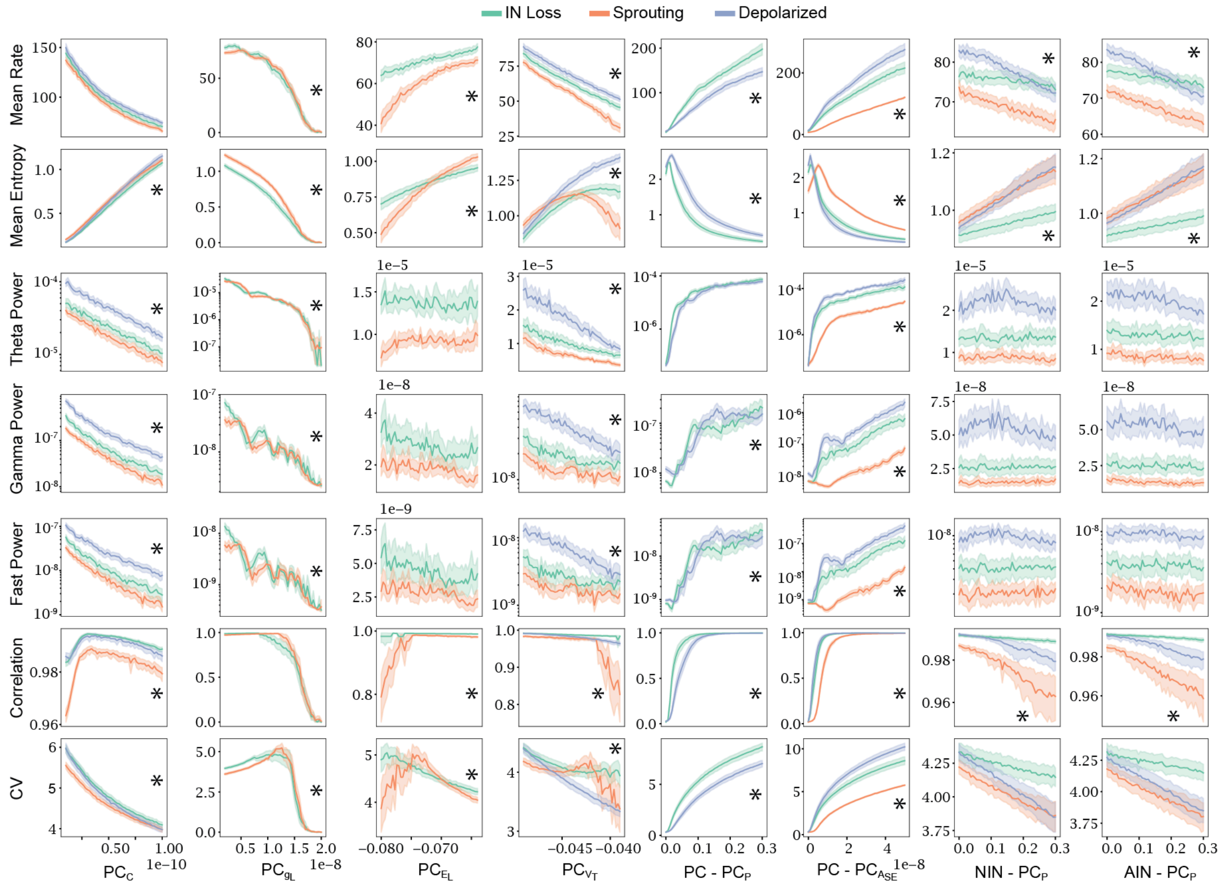


Figure Supp 6.1: An extended version of Figure 5 showing the effect of more parameters.

Table Supp 6.1: p-values of the ANOVA interaction between the parameter and the outcome.

| | PR(>F) | PR(>F) | PR(>F) | PR(>F) | PR(>F) | PR(>F) | PR(>F) |
|---|---|---|---|---|---|---|---|
| outcome | CV | Correlation | FastPower | GammaPower | MeanEntropy | MeanRate | ThetaPower |
| C(condition):C(AINPC_P) | 0.049790078 | 4.68E-21 | 0.999990844 | 0.999961673 | 1.80E-07 | 4.99E-13 | 0.996960627 |
| C(condition):C(NINPC_P) | 0.009287499 | 1.23E-23 | 0.604249686 | 0.509422927 | 1.03E-07 | 3.56E-10 | 0.879294295 |
| C(condition):C(PCPC_A_SE) | 2.73E-179 | 0 | 2.45E-212 | 1.34E-214 | 0 | 8.81E-274 | 2.54E-203 |
| C(condition):C(PCPC_P) | 6.22E-10 | 7.75E-284 | 2.69E-12 | 1.26E-12 | 1.95E-83 | 8.58E-25 | 0.036969746 |
| C(condition):C(PC_C) | 5.14E-13 | 6.00E-91 | 1.48E-236 | 0 | 8.82E-07 | 1 | 2.08E-156 |
| C(condition):C(PC_E_L) | 8.24E-26 | 2.52E-59 | 0.859895179 | 0.832443386 | 2.88E-95 | 1.20E-64 | 0.633548238 |
| C(condition):C(PC_V_T) | 2.80E-52 | 1.08E-109 | 5.23E-79 | 2.67E-78 | 0 | 5.09E-12 | 8.56E-57 |
| C(condition):C(PC_g_L) | 5.96E-10 | 0.001796825 | 1.47E-86 | 3.92E-86 | 5.78E-72 | 3.45E-08 | 2.11E-70 |
